# V-ATPase Disassembly at the Yeast Lysosome-Like Vacuole Is a Phenotypic Driver of Lysosome Dysfunction in Replicative Aging

**DOI:** 10.1101/2024.07.23.604825

**Authors:** Fiza Hashmi, Patricia M. Kane

## Abstract

Declines in lysosomal acidification and function with aging are observed in organisms ranging from yeast to humans. V-ATPases play a central role in organelle acidification and V-ATPase activity is regulated by reversible disassembly in many different settings. Using the yeast *Saccharomyces cerevisiae* as a replicative aging model, we demonstrate that V-ATPases disassemble into their V1 and V0 subcomplexes in aging cells, with release of V1 subunit C (Vma5) from the lysosome-like vacuole into the cytosol. Disassembly is observed after ≥5 cell divisions and results in overall vacuole alkalinization. Caloric restriction, an established mechanism for reversing many age-related outcomes, prevents V-ATPase disassembly in older cells and preserves vacuolar pH homeostasis. Reversible disassembly is controlled in part by the activity of two opposing and conserved factors, the RAVE complex and Oxr1. The RAVE complex promotes V-ATPase assembly and a *rav1Δ* mutant shortens replicative lifespan; Oxr1 promotes disassembly and an *oxr1Δ* mutation extends lifespan. Importantly, the level of Rav2, a subunit of the RAVE complex, declines in aged cells, and Rav2 overexpression delays V-ATPase disassembly with age. These data indicate that reduced V-ATPase assembly contributes to the loss of lysosome acidification with age, which affects replicative lifespan.

## INTRODUCTION

Lysosomal acidification is compromised with age in many organisms (Nixon, 2020). Reduced acidification has multiple downstream functional consequences. Lysosomal hydrolases operate optimally at the acidic pH, so cargo degradation is compromised at higher pH (Vilchez, Saez, & Dillin, 2014). Lysosomes are terminal compartments for autophagy pathways, so clearance of autophagic cargoes and recycling of nutrients, both critical in aging cells, is slowed (Hansen, Rubinsztein, & Walker, 2018; Kaushik et al., 2021). Iron and other heavy metals are sequestered in the acidic lysosomes; loss of sequestration can induce both oxidative stress (Diab & Kane, 2013) and deficiency in mitochondrial iron-sulfur proteins (Chen et al., 2020). Reduced lysosomal storage can also create toxic imbalances in amino acids such as cysteine that contribute to loss of mitochondrial function (C. E. Hughes et al., 2020). Recent work highlights the central role of the lysosome in nutritional signaling; many aspects of this signaling are linked to acidification (Perera & Zoncu, 2016). Loss of lysosomal acidification impacts many processes associated with age-related functional decline, but the mechanisms behind increased lysosomal pH are not fully understood.

The highly conserved vacuolar H^+^- ATPase (V-ATPase) acidifies lumens of lysosomes and lysosome-like vacuoles, as well as endosomes and the late Golgi apparatus (Collins & Forgac, 2020). V-ATPases are multi-subunit protein complexes that couple ATP hydrolysis to proton pumping into organelle lumens. The V-ATPase consists of two subcomplexes: a peripheral V1 subcomplex that is responsible for ATP hydrolysis connected to a membrane- embedded V0 subcomplex containing the proton pore. V-ATPase subunit sequences are conserved across eukaryotes and recent V-ATPase structures indicate very strong structural similarity between yeast and mammalian V-ATPases (Oot & Wilkens, 2020).

V-ATPase activity is highly regulated and responsive to environmental conditions. Reversible disassembly is a versatile mechanism of V-ATPase regulation that fine-tunes the activity of the proton pump to meet cellular demands (Collins & Forgac, 2020; Jaskolka, Winkley, & Kane, 2021). In reversible disassembly, the V1 subcomplex is released from the V0 subcomplex inhibiting both ATP hydrolysis and proton pumping (Kane, 1995; Sumner et al., 1995). V1 subunit C dissociates from both subcomplexes and becomes cytosolic during disassembly (Kane, 1995). Disassembly is post-translational and rapidly reversible (Kane, 1995). It was first observed in the yeast *S. cerevisiae* and the tobacco hornworm *M. sexta* upon acute glucose deprivation and was reversed by glucose replenishment (Kane, 1995; Sumner et al., 1995). It has since become clear that reversible disassembly occurs widely and in response to diverse signals. For example, many cells appear to promote V-ATPase reassembly under conditions of nutrient deprivation and mTOR inhibition, possibly as a means of promoting lysosomal proteolysis and nutrient recycling (Ratto et al., 2022). In neurons, V-ATPases are reversibly disassembled as part of each synaptic vesicle cycle (Bodzeta, Kahms, & Klingauf, 2017). In cardiomyocytes, lipid overload can promote V-ATPase disassembly in endosomes, ultimately contributing to long-term insulin resistance (Liu et al., 2017). Reversible disassembly can also be manipulated by both host cells and pathogens to prevent or facilitate infection (Kohio & Adamson, 2013).

The RAVE (Regulator of Acidification of Vacuoles and Endosomes) complex and TLDc (Tre2/Bub2/Cdc16 LysM domain catalytic) protein Oxr1 regulate reversible disassembly of the V-ATPase (Jaskolka, Winkley, et al., 2021; Khan et al., 2022; Klossel et al., 2024; Seol, Shevchenko, & Deshaies, 2001). The RAVE complex consists of three subunits: Rav1, Rav2, and Skp1 and is required for reassembly of V-ATPase complexes disassembled by glucose deprivation. In mutants lacking Rav1 or Rav2, V-ATPases are disassembled into V1 and V0 subcomplexes and vacuolar acidification is lost (Jaskolka, Winkley, et al., 2021). The role of Skp1 in RAVE is less clear and it may be dispensable for RAVE function (Jaskolka, Winkley, et al., 2021). The RAVE complex associates with V1 subcomplexes in the cytosol (Jaskolka, Tarsio, Smardon, Khan, & Kane, 2021). In contrast, recent studies suggest that Oxr1, a member of the TLDc family of proteins, promotes disassembly of V-ATPases. Mutants lacking Oxr1 maintain higher levels of assembled V-ATPases in the absence of glucose (Khan et al., 2022; Khan & Wilkens, 2024; Klossel et al., 2024).

Several mechanisms for loss of lysosomal acidification with age have been proposed. In yeast, the long-lived plasma membrane proton pump, Pma1, accumulates in mother cells, and it has been proposed that increased proton export through Pma1 disrupts the balance of Pma1 and V-ATPase activities and compromises organelle acidification (Devare et al., 2020; Henderson, Hughes, & Gottschling, 2014). Early loss of vacuolar acidification in yeast has been correlated with defective mitochondrial morphology and function. Both vacuolar deacidification and mitochondrial morphology phenotypes are suppressed by overexpression of the V-ATPase catalytic subunit, *VMA1*, or an ER-localized assembly factor, *VPH2 (VMA12)*, suggesting a possible deficiency in these factors with age (A. L. Hughes & Gottschling, 2012). In *C. elegans*, stability of the *VMA1* transcript is controlled by a microRNA, miR-1, which can globally control lysosomal acidification (Schiffer et al., 2021). Wilms et al. (Wilms et al., 2017) demonstrated that deletion of the mTOR effector Sch9 promotes V-ATPase assembly and extends yeast chronological lifespan (CLS). All of these mechanisms could contribute to loss of vacuolar/lysosomal acidification.

However, despite the importance of reversible disassembly in V-ATPase regulation, V-ATPase assembly state during aging has not been explored extensively. Here we show that V-ATPase assembly changes with age in a yeast replicative aging model. Increased V-ATPase disassembly in older cells is accompanied by decreased vacuolar acidification that does not appear to stem from reduced V-ATPase subunit levels. Instead, we provide evidence that reduced activity of the RAVE (Regulator of H^+^-ATPase of Vacuolar and Endosomal membranes) assembly complex may give rise to net V-ATPase disassembly and increased lysosomal pH with age.

## RESULTS

### V-ATPases are more disassembled and vacuoles more alkaline after ∼5 cell divisions

Given the evidence that vacuoles and lysosomes are less acidic in older cells, we hypothesized that V-ATPase assembly and activity might also change with age. Replicative aging in the yeast *S. cerevisiae* is a widely accepted model for aging (He, Zhou, & Kennedy, 2018).

Yeast cells divide asymmetrically with each cell division giving rise to a new, “rejuvenated” daughter cell from an established mother cell. Each cell division leaves a bud scar on the mother, allowing visual assessment of age. The yeast V1C subunit (Vma5) is released from both V1 and V0 during V-ATPase disassembly, so we first visualized Vma5-GFP localization in a mixed age population of cells (Figure 1). (Vacuoles are visible as indentations under DIC (differential interference contrast) optics.) In parallel, we monitored replicative age of each cell by staining with calcofluor white (CW) which labels the bud scars on mother cells. As shown in Figure 1a, Vma5-GFP is tightly localized to the vacuolar membrane in cells with few or no bud scars. In contrast, in older cells with more bud scars, Vma5-GFP exhibited a marked decrease in fluorescence at the vacuolar membrane and a notable increase in cytosolic fluorescence. To compare Vma5-GFP localization between cells, we quantitated maximum fluorescence from a line scan across each cell. As shown in Figure 1b, a line scan from a young cell has prominent peaks corresponding to edges of the vacuole with a high maximum fluorescence signal, while older cells have less prominent peaks. We conducted the same analysis across populations of cells of mixed age, normalized to the maximum fluorescence signal of young cells and binned the results by the number of bud scars. As shown in Figure 1c, young cells, defined as having less than 5 bud scars, displayed Vma5-GFP localization at the vacuolar membrane. However, the normalized maximum fluorescence signal, representative of vacuole localization, decreases significantly in cells with five bud scars or more. This early onset of V-ATPase disassembly aligns with previous reports indicating declines in lysosomal pH early in replicative aging (A. L. Hughes & Gottschling, 2012).

**Figure 1:**
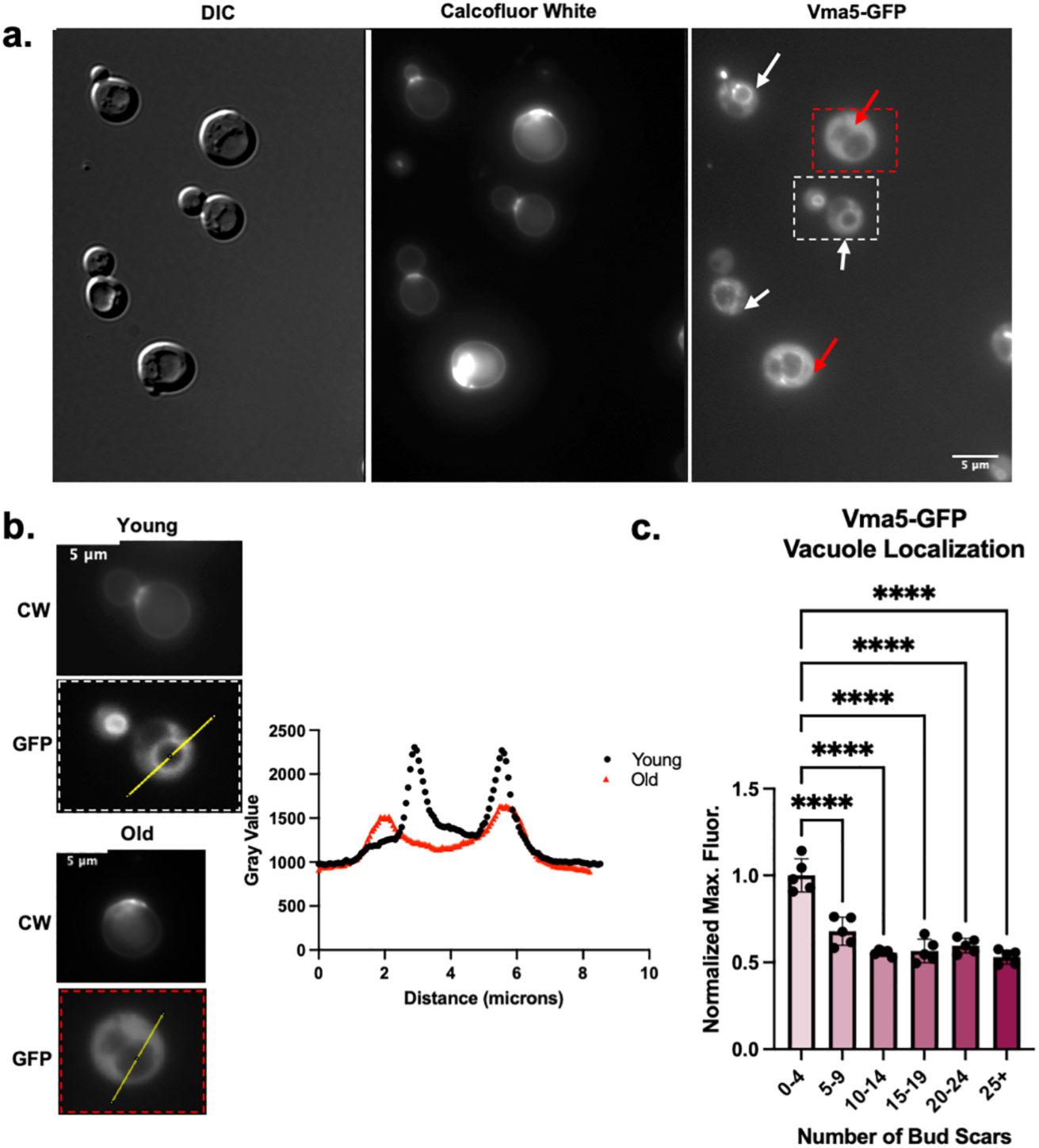
V-ATPases are more disassembled in yeast cells of older replicative age. **(a)** Vma5- GFP cells grown in SC containing 2% glucose. DIC was used to visualize vacuoles. Bud scars are stained with CW to determine replicative age. **(b)** Left panel provides representative example of quantitative measurements using line scans. The young and old cells are from the dashed white and red boxes, respectively, in the Vma5-GFP image of **Figure 1a**. Plot profiles are superimposed for the young (black) and old (red) cells. **(c)** Quantitation of maximum fluorescent intensity across five biological replicates, after normalization to the average intensity in the youngest bin for each replicate. Each biological replicate (dot) represents at least 20 cells and bars represent the mean +/- s.e.m. “Old” for subsequent experiments (without age enrichment) is categorized as ≥5 bud scars. **** represents a p-value <0.0001 calculated by ordinary one-way ANOVA.

Vma5 is a V1 subunit that bridges the V1 and V0 subcomplexes of the V-ATPase. The V1 subcomplex also contains Vma2, and the V0 subcomplex contains Vph1, which comprises part of the proton pore (Figure 2a). Vma5 appears to be released most completely from the membrane by reversible disassembly (Tabke et al., 2014), but the rest of V1 also dissociates from the vacuolar membrane. We assessed the cellular distribution of Vma2-GFP and Vph1-GFP. There was less membrane-bound Vma2-GFP in older cells relative to younger cells (Figure 2b), when assessed as in Figure 1. However, vacuolar Vph1-GFP levels were the same between old and young cells (Figure 2c). These results are consistent with disassembly of the V-ATPase as cells age.

**Figure 2:**
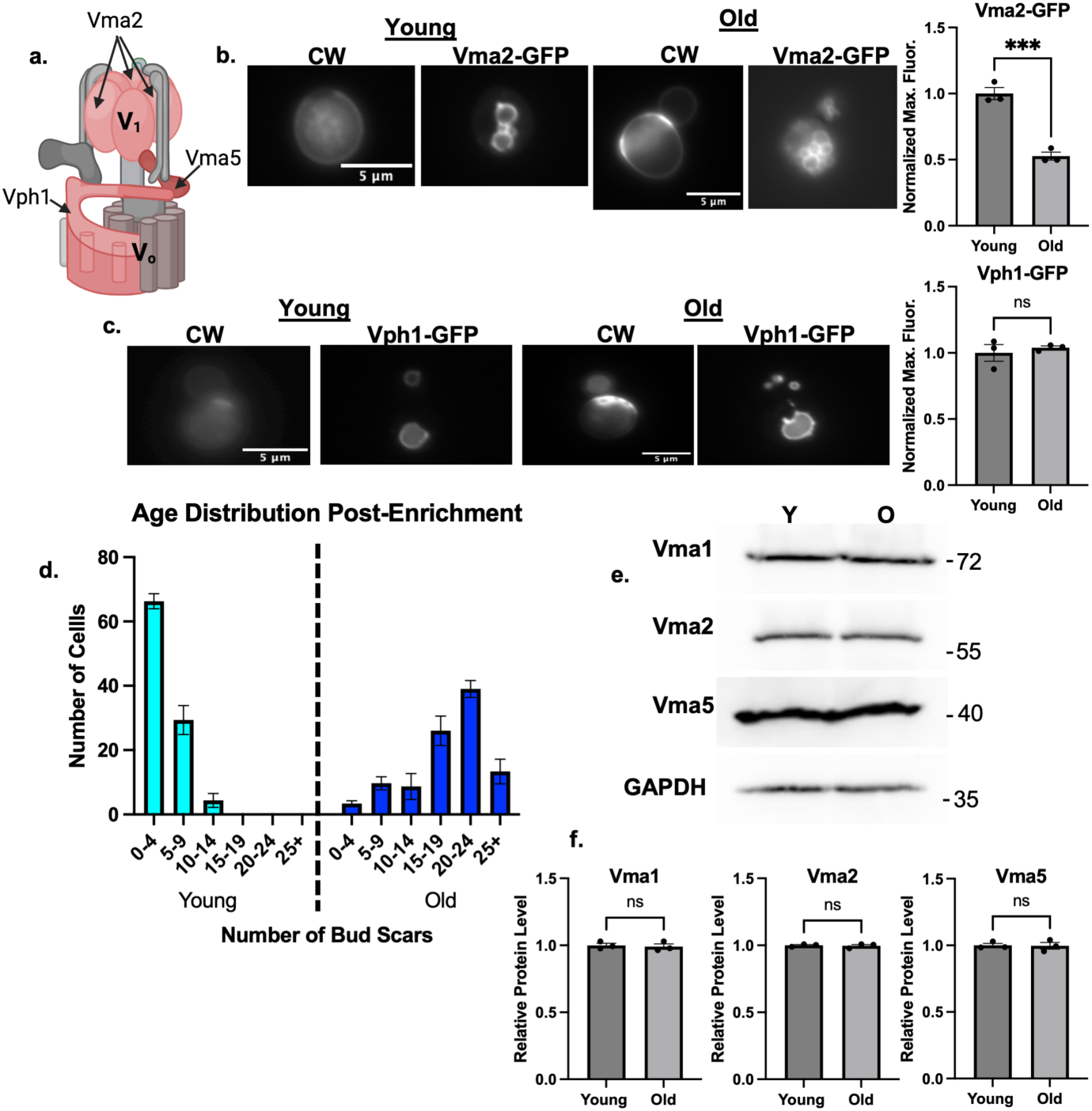
Age-enriched populations of cells have comparable levels of V-ATPase subunits. (a) Diagram of V-ATPase showing the positions of Vma5, Vma2, and Vph1. (b) Vma2-GFP cells (V1B) grown in SC with 2% glucose. CW was used to visualize bud scars. The CW images for young cells were overexposed relative to those for the old cells in order to visualize the low level staining in young cells. Normalized maximum fluorescence was obtained as in Figure 1. Means +/- s.e.m. of three biological replicates are shown; each replicate is represented by a Significance was assessed by unpaired Student’s *t* test, *** p value=0.0009 . **(c)** Vph1-GFP cells were visualized and analyzed as described in **2b**. **(d)** After biotin-streptavidin age enrichment, bud scars were counted for 100 cells in young and old populations, then binned by bud count. **(e)** Lysates from age-enriched populations defined as in **1c** were separated by SDS-PAGE. Immunoblots for the indicated proteins in young (Y) vs. old (O) cell populations as defined as in 1c. (f) Band intensities were quantified using FIJI, ratios of subunit levels to the GAPDH control were calculated and normalized to the young population for each biological replicate. Data are presented as mean (horizontal bars) ± s.e.m. (whiskers) of three biological replicates. n.s.= not significant by unpaired Student’s *t* test.

Although comparable levels of Vph1 at the vacuole suggests that expression of V0 subunits and V0 assembly are intact, V1 subunits could become cytosolic because of reduced V1 subunit levels in older cells. To address this question, we isolated populations old and young yeast cells by biotinylating the cell walls in a mixed age population, allowing growth to continue for several generations, and then obtaining “old” cells by biotin-streptavidin affinity chromatography (Jin, Cao, & Liu, 2021). Daughter cells that emerged after biotinylation cannot bind to magnetic streptavidin beads and represent the “young” population. The age distribution was determined by counting bud scars in each population and binning by the number of buds per cell (Figure 2d). When prepared by this method, the population of old cells peaks at 20-24 bud scars, while there was a median value of 0-4 bud scars in the young population. Cell lysates were prepared from each population and analyzed by SDS-PAGE and immunoblotting. As shown in Figures 2e and f, there is no significant difference in the cellular levels of V1 subunits Vma1, Vma2, and Vma5 between young and old populations. These results indicate that the cytosolic populations of V1 subunits arise from disassembly of the V-ATPase, rather than inability to assemble because of lack of V-ATPase subunits.

Reversible disassembly of V-ATPases is employed in a number of contexts to provide dynamic regulation of the complex in response to changing cellular conditions. Disassembled V1 and V0 subcomplexes lack ATPase and proton transport activity and ATP-driven proton pumping is restored upon reassembly. In order to test whether the lower levels of assembled V-ATPases in old cells result in reduced capacity for acidification, we measured the response of young and old cells to an acute glucose deprivation (an abrupt shift to 0% glucose), which promotes disassembly of the yeast V-ATPase, followed by readdition of glucose to a 2% final concentration, which promotes reassembly and reactivation of the complex (Kane, 1995).

Young and old yeast cells obtained as described above were loaded with the ratiometric pH sensor BCECF-AM (2’,7’-Bis-(2-carboxyethyl)-5-(and-6)-carboxyfluorescein acetoxy methyl ester) which localizes to the vacuole in yeast cells (Diakov, Tarsio, & Kane, 2013). Both young and old cells were shifted to medium with no glucose for ∼30 minutes. Fluorescence of cell suspensions was then monitored continuously (Figure 3a) and glucose was added at the indicated time. This assay revealed clear age-dependent differences in the vacuolar pH response to glucose stimulation. Young cells exhibit a rapid drop in vacuolar pH upon glucose addition. This drop was previously shown to be V-ATPase-dependent and to correlate with V-ATPase assembly (Martinez-Munoz & Kane, 2008). In old cells, however, there was a smaller pH decrease after glucose addition, indicating a more alkaline vacuole and consistent with the lower levels of V-ATPase assembly observed by microscopy during growth in glucose-replete media in Figures 1 and 2. Quantitative analysis of pH at defined time points across biological replicates (Figure 3b) indicates that vacuoles in old cells are significantly more alkaline than their younger counterparts at each of the indicated time points. These results suggest an age-related alteration in vacuolar pH regulation through the inability of V-ATPase to reassemble. As a result, vacuoles in old cells display a more alkaline pH than vacuoles in young cells.

**Figure 3:**
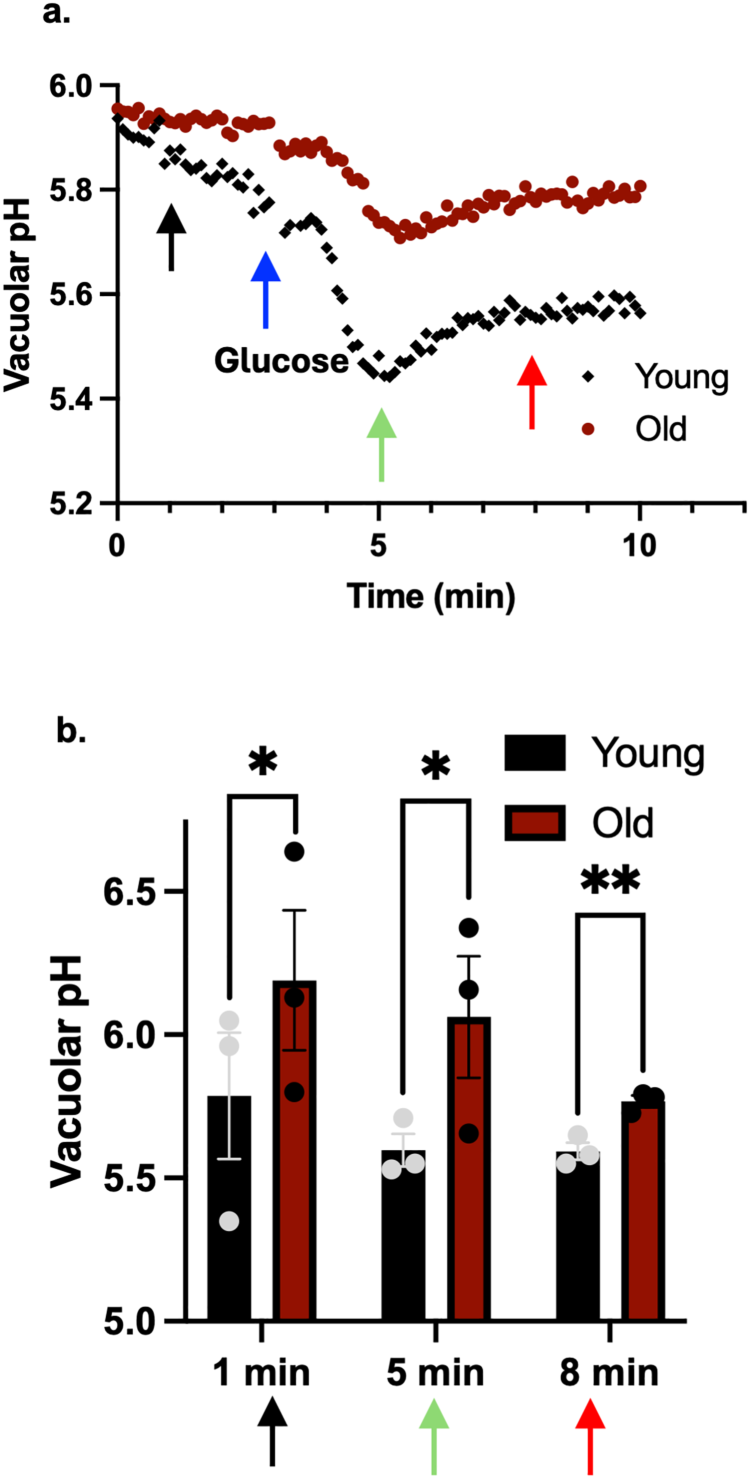
Vacuolar pH is more alkaline in old cells. (a) Vacuolar pH responses were measured for wild-type BY4742 age-enriched young and old populations as described in Methods. Glucose-deprived cultures were loaded with BCECF-AM. Fluorescence intensity values were collected every 10 sec and glucose was added to a final concentration of 2% after 3 min. The ratio of fluorescence signals was converted to pH as described in Methods. (b) pH measurements at 1 min (before glucose addition), 5 min (2 min after glucose addition), and 8 min (5 min after addition) are presented as mean (horizontal bars) ± s.e.m. (whiskers) of three biological replicates. * = p value of 0.02, ** p value <0.001.

### Caloric restriction restores V-ATPase assembly and vacuolar acidification in older cells

Caloric restriction (CR) is defined as a reduction in caloric intake in the presence of adequate nutrition (Longo & Anderson, 2022). CR promotes both cellular health and longevity. Extensive research in organisms ranging from yeast to worms, flies, and rodents suggests that CR can have significant anti-aging effects and promote overall health (Longo & Anderson, 2022). For *S. cerevisiae*, adjusting the concentration of glucose in the growth medium from the 2% used above to 0.5% is a common way to induce CR. This treatment does not significantly reduce growth rate over several cell divisions (Supporting information Figure 1a).

As shown in Figure 4, CR reverses the V-ATPase disassembly in older cells. Notably, the V1 subunits Vma5-GFP (Figure 4a) and Vma2-GFP (Figure 4b) are recruited to the vacuolar membrane in both young and old cells, and there is no significant difference in fluorescence at the vacuolar membrane with age. Fluorescence intensities of Vph1-GFP continue to be similar between old and young cells (Figure 4c). This suggests that CR extends V-ATPase assembly beyond the replicative age of ≥5 bud scars when cells grown in higher glucose begin to show disassembly.

**Figure 4:**
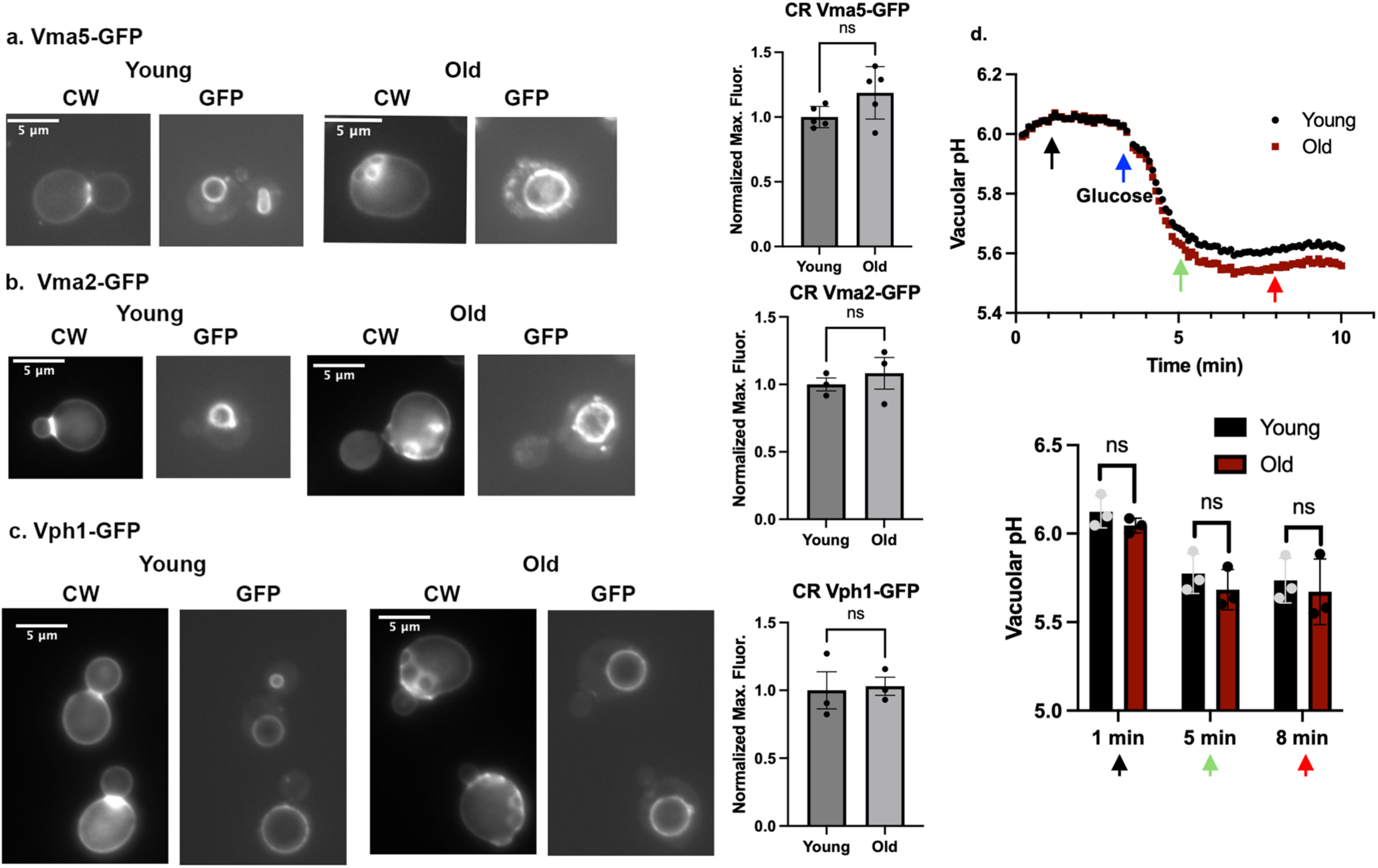
C**a**loric **restriction restores V-ATPase assembly and vacuolar pH in old cells. (a-c)** Strains used in **Figures 1** and **2** show recruitment of Vma5-GFP (**a)** and Vma2-GFP **(b)** to the vacuolar membrane after growth in SC with 0.5% glucose. Vph1-GFP (**c)** remains unchanged. Normalized maximum fluorescence was quantitated as described in Methods. No significant differences between young and old cells were observed by unpaired Student’s *t* test. Designation of young and old cells by CW staining was as described in **Figure 1**. **(d)** Young and old cell populations were obtained by biotin-streptavidin age-enrichment from cells grown in YEP supplemented with 0.5% glucose, and vacuolar pH responses were measured. Data were collected and analyzed as in **Figure 3b**. Data are presented as mean (horizontal bars) ± s.e.m. (whiskers) of three biological replicates.

We hypothesized that vacuolar pH in old cells might also be restored. We grew cells under CR conditions and monitored vacuolar pH before and after addition of glucose as described above. Figure 4d demonstrates that the glucose-stimulated decrease in vacuolar pH, which was compromised in older cells grown in 2% glucose (Figure 3), was restored to the level of young cells in cells after growth under CR conditions. This observation indicates that CR has a direct impact on both V-ATPase assembly and vacuolar acidification in aging cells. In addition, it further highlights the potential connection between V-ATPase assembly, vacuolar acidification and aging.

These results point to nutritional signaling as an important determinant of V-ATPase assembly and activity in aging cells. Bond and Forgac demonstrated that V-ATPase assembly was preserved under low glucose conditions in an *ira2Δ* mutant. Deletion of *IRA2*, or its paralogue *IRA1*, activates the Ras/protein kinase A (PKA) pathway (Bond & Forgac, 2008). We hypothesized that V-ATPase assembly might be preserved in older cells, mimicking the effects of CR, in an *ira2Δ* mutant. We visualized Vma5-GFP localization and as shown in Supporting Information Figure 1b, the *ira2Δ* cells do maintain V-ATPase assembly in older cells. Therefore, we determined replicative lifespan (RLS) for the *ira2Δ* mutant by daughter dissection (Supporting Information Figure 1c). The median lifespan of the *ira2Δ* mutant is significantly shorter than that of wild-type cells. These data are consistent with evidence that upregulation of the Ras/protein kinase A pathway shortens lifespan (Longo & Anderson, 2022) and suggest that although there is improved V-ATPase assembly in older cells in the *ira2Δ* mutant, it is not sufficient to balance effects on other targets of the pathway in aging cells.

### Regulators of V-ATPase assembly affect replicative lifespan

We next sought to manipulate V-ATPase assembly more directly in order to assess the significance of maintaining V-ATPase assembly and activity in aging cells. We hypothesized that the RAVE complex and Oxr1, cellular factors that promote V-ATPase assembly and disassembly as shown in Figure 5a, might also affect RLS.

**Figure 5:**
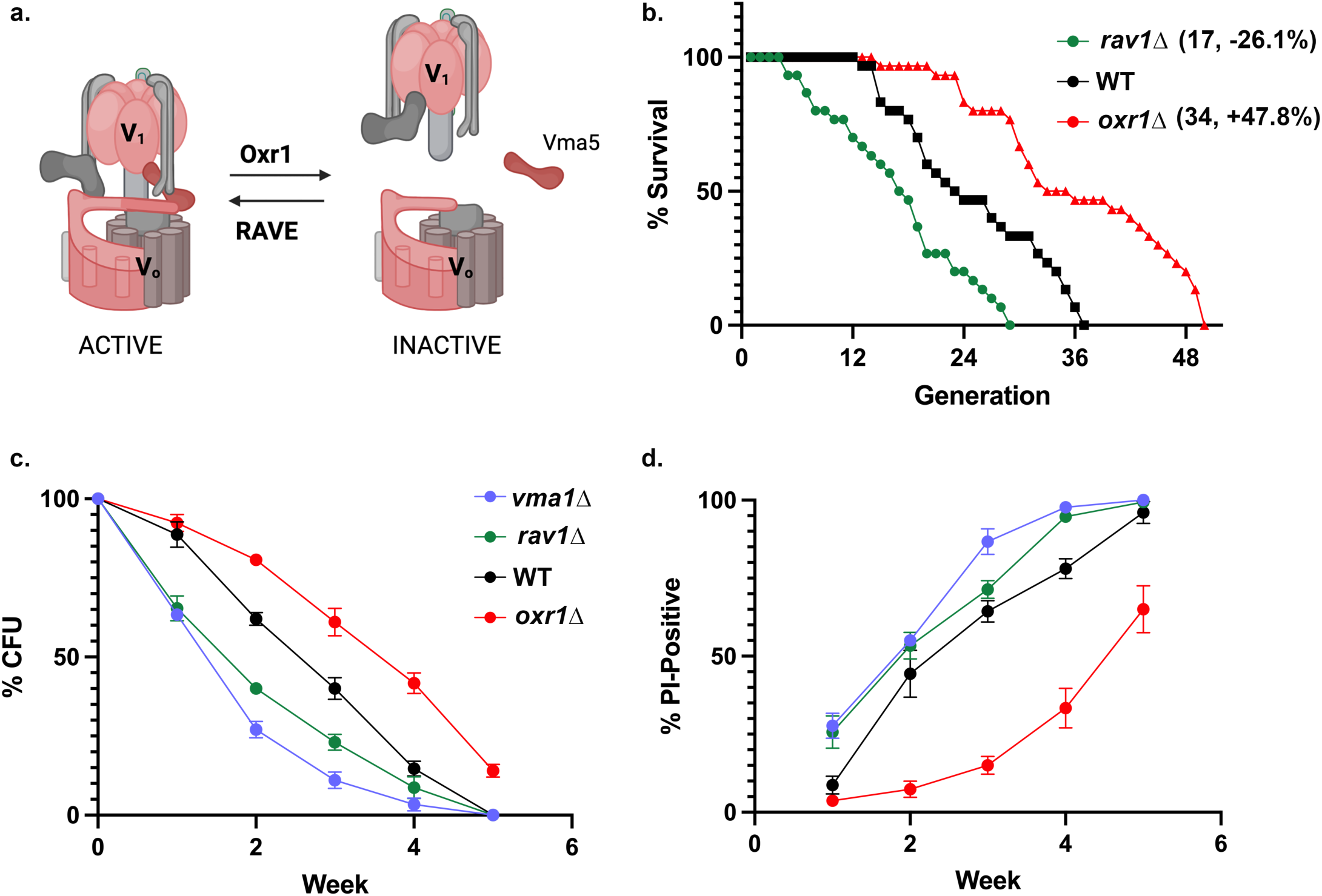
E**f**fects **of *rav1Δ* and *oxr1Δ* on RLS and CLS. (a)** Schematic of reversible disassembly. **(b)** Kaplan-Meier curves comparing the RLS of *rav1Δ* (green), *oxr1Δ* (red), and wild-type cells (black). Median number of replicative generations is shown in parentheses, with the % difference between median values for *rav1Δ* and *oxr1Δ* vs. wild-type. Deletion of *OXR1* increases RLS (p<0.001), and deletion of RAVE component *RAV1* shortens RLS (p < 0.01). **(c, d)** Plots of colony-forming units **(c)** and percentage of cells stained with propidium iodide **(d)** vs. time. Each point is the mean of three biological replicates. Strains included are defined to the right of **(c)**. Statistical analysis of **(c)** indicates a p-value <0.0001 for all mutants in comparison to wild-type. In **(d)**, *vma1Δ* and *oxr1Δ* differed from wild-type with a p-value <0.0001, and *rav1Δ* with a p-value= 0.0007.

To explore the functional significance of these assembly regulators in aging cells, we examined the RLS of deletion mutants lacking Rav1 or Oxr1 (Figure 5b). RLS was measured at pH 5, conditions that are optimal for growth of *rav1Δ* strains. Deletion of Rav1 shortened RLS (median 17 cell divisions, n= 30) by 26% relative to wild-type cells (median 23 cell divisions, n= 30). In contrast, the *oxr1Δ* mutation extended RLS by 47.8% over wild-type (median 34 cell divisions, n=30). These results reinforce the significance of V-ATPase assembly in RLS and suggest that the RAVE pro-assembly activity (disrupted in the *rav1Δ* mutant) and Oxr1 anti- assembly activity (lost in *oxr1Δ*) may be central determinants of lifespan.

We next asked whether loss of RAVE and Oxr1 affects chronological aging of yeast cells. Replicative aging assesses the number of times a mother cells can divide, while chronological aging assesses cell survival as nutrients are depleted over time. Wild-type cells and *rav1Δ*, *oxr1Δ*, and *vma1Δ* mutants were grown to saturation in medium buffered to pH 5.5 (Murakami et al., 2012; Wilms et al., 2017). (*VMA1* encodes the catalytic subunit of the V-ATPase and the *vma1Δ* mutant has no V-ATPase activity.). At the indicated times, aliquots were removed from each culture, cell density was determined, and a defined number of cells was plated on fresh medium. The percentage of each cell type that formed colonies (c.f.u.) on fresh plates at each time is shown in Figure 5c. Even in buffered medium, which improves the survival of *vma* mutants (Wilms et al., 2017), less than half of the *vma1Δ* and *rav1Δ* mutant cells are able to form colonies after 2 weeks. In contrast, almost half of the *oxr1Δ* cells form colonies at 4 weeks, and wild-type cells have an intermediate phenotype. The percentage of cells staining with propidium iodide, an indication of cell permeabilization and death, increases as colony formation declines (Figure 5d). However, it is notable that even after 5 weeks, only ∼60% of *oxr1Δ* cells stain with propidium iodide. These results complement the replicative aging data and indicate that mutations that preserve V-ATPase assembly can significantly increase lifespan, while mutations that compromise activity and/or assembly decrease lifespan. Yeast encode a second isoform of the V0 a-subunit, Stv1, which localizes to the Golgi in wild-type cells. Stv1-containing V-ATPases do not require RAVE for assembly (Smardon et al., 2014), and Stv1 has been shown to relocalize to the vacuole in an *oxr1Δ* mutant (Klossel et al., 2024). In order to test whether the lifespan extension in *oxr1Δ* cells arises from Stv1-containing V-ATPases at the vacuole, we constructed a *stv1Δoxr1Δ* double mutant strain and assessed its RLS and V-ATPase assembly. As shown in Supporting Information Figure 2a, the double mutant strain retains the long lifespan of the *oxr1Δ* relative to wild-type, suggesting that vacuolar Stv1 is not responsible for this effect.

Rather, V-ATPases containing the Vph1 isoform account for the assembly in older cells of the *oxr1Δ* mutant (Supplemental Information Figure 2b).

Given the results with these deletion mutants, we asked whether there were differences in levels of the RAVE subunits or Oxr1 between young and old cells. We isolated young and old populations of cells containing functional myc13-tagged Rav1 or FLAG-tagged Rav2 by biotinylation and streptavidin magnetic separation as described above (Figure 2), then assessed the levels of the tagged proteins in cell lysates. Although there is no significant difference in Rav1-myc13 levels between young and old cells (Figure 6a), we consistently observed a significant decrease in the level of Rav2-FLAG in older cells (Figure 6b). We assessed expression of *RAV2* in young and old cells by quantitative PCR but observed no difference in mRNA levels (Supporting Information Figure 2c). Because Rav2 is required for RAVE complex function in promoting V-ATPase assembly (Seol et al., 2001; Smardon, Tarsio, & Kane, 2002), these results suggest that partial loss of RAVE function in old cells could contribute to reduced

**Figure 6:**
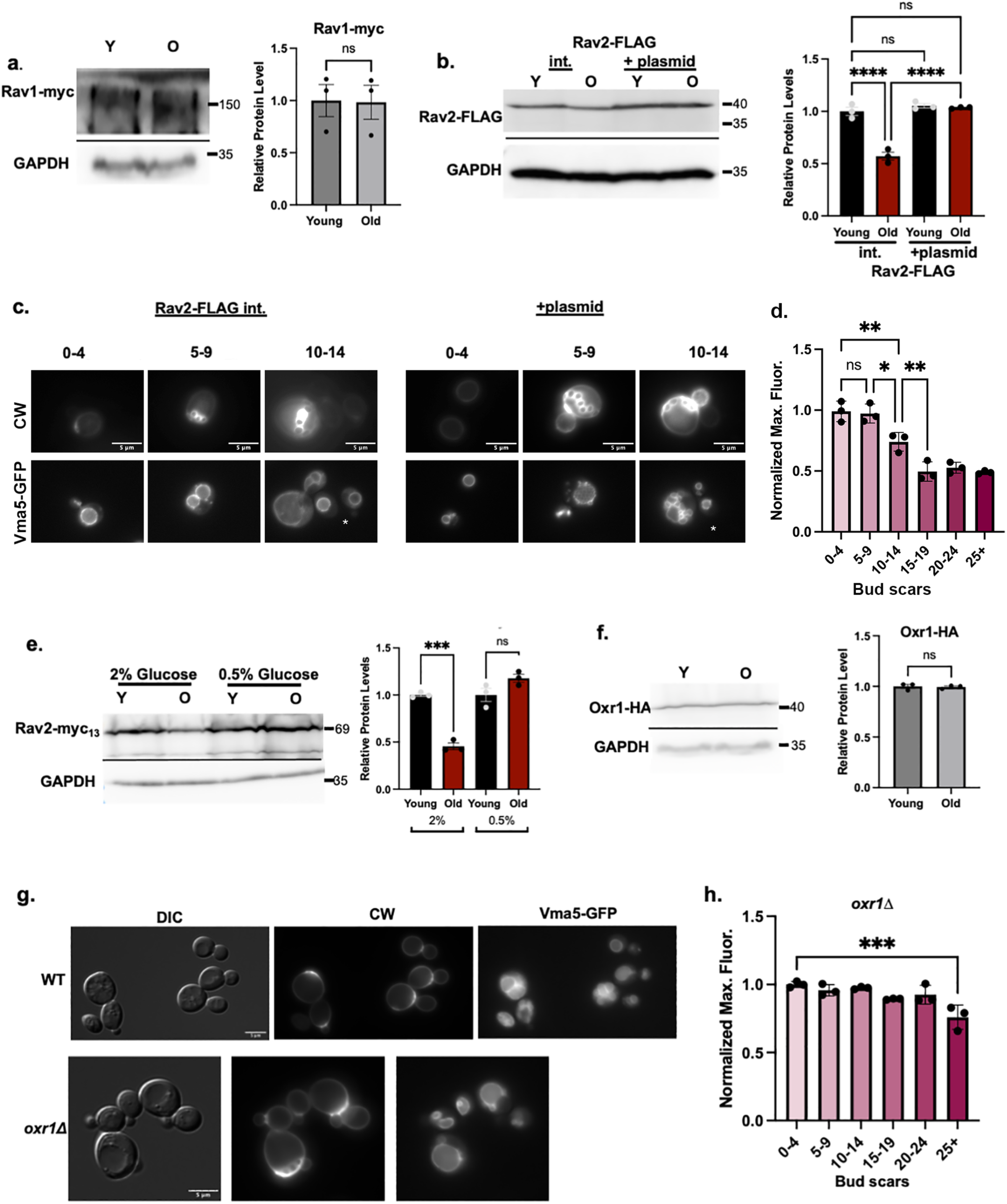
R**A**VE **subunit and Oxr1 levels in aging cells. (a)** Rav1 levels in young and old cells. Age-enriched populations were obtained from WT cells containing Rav1-myc13 (young (Y) versus old (O) cells). Cell lysates were prepared and analyzed as in **Figure 2**. Normalized data are presented as mean (horizontal bars) ± s.e.m. (whiskers) of three biological replicates. ns=not significant. **(b)** Rav2-FLAG levels from young and old cells prepared as in **6a** from haploid cells containing only a chromosomally integrated (int.) Rav2-FLAG or cells with both integrated Rav2-FLAG and low copy Rav2-FLAG plasmid (+plasmid). Normalized data are presented as mean (horizontal bars) ± s.e.m. (whiskers) of three biological replicates. ****, p-value<0.0001, ns= not significant. **(c)** Rav2-FLAG Vma5-GFP cells, without (int.) or with the Rav2-FLAG plasmid (+ plasmid). CW is used to visualize bud scars as in **Figure 1**; white stars on the Vma5- GFP images indicate newborn cells with no bud scars. Images representative of cells with 0-4, 5-9, and 10-14 bud scars are shown. **(d)** Rav2-FLAG cells + plasmid were binned by the number of bud scars and normalized maximal fluorescence quantitated as in **Figure 1**. **, p-value<0.01 and *, p-value<0.05 for the indicated comparisons; n.s.=not significant. **(e)** Immunoblot analysis comparing Rav2-myc13 levels in cells grown in YEP with 2% glucose or 0.5% glucose (CR conditions). Samples were prepared, analyzed, and presented as in **6a**. ***, p- value <0.001, n.s.= not significant. **(f)** Immunoblot of lysates from *oxr1Δ* strain expressing Oxr1-HA from a low copy plasmid. Young and old cells were isolated as in **6a**. Data are presented as mean (horizontal bars) ± s.e.m. (whiskers) of three biological replicates. **(g)** Wild-type and *oxr1Δ* cells containing Vma5-GFP grown in SC with 2% glucose. DIC and CW were used to visualize vacuoles and bud scars as in **Figure 2b**. **(h)** *oxr1Δ* cells were binned by the number of bud scars and normalized maximal fluorescence quantitated as in **Figure 1**. ***, p-value <0.001 in comparison to the youngest bin; there is no other significant difference.

V-ATPase assembly and replicative aging. We transformed the cells containing a chromosomal copy of Rav2-FLAG with an additional copy of Rav2-FLAG on a low-copy plasmid to determine whether we could increase Rav2 levels and improve V-ATPase assembly in older cells. As shown in Figure 6b, addition of the plasmid increases Rav2-FLAG levels in old cells. We compared Vma5-GFP localization with and without the Rav2-FLAG plasmid in a mixed age population in Figure 6c. In cells overexpressing Rav2-FLAG, Vma5-GFP is at higher levels in the vacuoles of cells with ≥5 bud scars. The quantitation and binning by bud scar number in

Figure 6d confirms that there is no significant decline in vacuolar Vma5-GFP in cells with 5-9 bud scars and only in cells with ≥ 15 bud scars does vacuolar fluorescence reach the level in “old” wild-type cells (Figure 1). These data indicate that increasing Rav2-FLAG levels can help preserve V-ATPase assembly in older cells.

We hypothesized that restoration of V-ATPase assembly in older cells grown under CR conditions (Figure 4) might be supported by restoration of Rav2 levels. To test this, we isolated young and old populations from Rav2-myc13 tagged cells grown under CR conditions. Under these conditions, V-ATPase assembly and function is restored in older cells (Figure 4) and Rav2 levels are also restored (Figure 6e).

Alterations in Oxr1 levels with age could also affect V-ATPase assembly. We isolated young and old populations from a strain containing HA-tagged Oxr1 and found that the levels of Oxr1-HA in whole cell lysates do not change with age (Figure 6f). To further explore the basis of the extended RLS in *oxr1Δ* cells, we observed Vma5-GFP localization in a population of *oxr1Δ* cells of mixed age. As shown in Figure 6g, h, *oxr1Δ* mutants localize Vma5-GFP to the vacuole even in older cells with ≥5 bud scars. To quantitate this effect, we again binned cells by the number of bud scars as described above, and compared Vma5-GFP localization to localization in young daughter cells (Figure 6h). In contrast to wild-type cells, there is no significant difference in Vma5-GFP localization in the *oxr1Δ* mutant until cells have divided 25 or more times.

Intriguingly, these results suggest that the *oxr1Δ* mutation mirrors the effects of CR on V-ATPase assembly and longevity. The results also indicate that direct manipulation of V-ATPase assembly by mutation of critical assembly factors can affect replicative and chronological lifespan, both of which are linked to cellular pH homeostasis (Murakami et al., 2012).

Specifically, promoting V-ATPase assembly through the RAVE complex appears to be important for preserving lifespan, while the opposing effects of Oxr1 on the V-ATPase tend to shorten lifespan.

## DISCUSSION

These results establish V-ATPase disassembly as an important factor in reduced vacuolar acidification in aging cells. We demonstrate an increase in V-ATPase disassembly at relatively early replicative ages (5-9 cell divisions) similar to the age at which Hughes and Gottschling first observed compromised vacuolar acidification (A. L. Hughes & Gottschling, 2012). We show that promoting assembly of the V-ATPase through CR can restore V-ATPase assembly and vacuolar pH in older cells. Deletion of Oxr1, a negative regulator of V-ATPase assembly, can extend both replicative and chronological lifespan. In contrast, mutation of the RAVE complex shortens lifespan and levels of the Rav2 subunit decline with age. Taken together, these data support V-ATPase disassembly as a significant age-related factor behind reduced vacuolar and lysosomal acidification and the associated declines in function.

This mechanism does not necessarily conflict with those proposed previously. If V-ATPases are more disassembled in older cells, cells may be less able to tolerate an imbalance between Pma1 activity at the plasma membrane and V-ATPase activity at the vacuole (Henderson et al., 2014). Okreglak et al. (Okreglak et al., 2023) recently visualized oscillations of vacuolar pH in mother cells that were coordinated with cell cycle. Oscillations cease upon V-ATPase inhibition, but it is not clear whether reversible disassembly of the V-ATPase is required. It is possible that this oscillatory behavior is lost in older mother cells, leading to loss of their ability to restore vacuolar pH after cell division. Notably, we see no difference in protein levels of V1 subunits Vma1, Vma2, and Vma5, between young and old cells. The similar levels of Vph1 in vacuoles of young and old cells suggest that V0 subunits are present at similar levels, since V0 assembly occurs in the ER and reduced levels of any V0 subunit reduces V0 subcomplex levels at the vacuole. The data presented here suggest a post-transcriptional regulatory mechanism, but increased levels of a biosynthetic assembly factor (A. L. Hughes & Gottschling, 2012) could promote assembly and improve acidification.

Many questions remain to be investigated. The reduction in Rav2 protein levels with age appears to be post-transcriptional but we do not yet know whether this arises from reduced translation or increased degradation. It is intriguing in young cells, addition of an extra copy of Rav2-FLAG did not increase levels of the protein in Figure 6b, suggesting the Rav2 subunit levels are controlled. Perhaps more importantly, even though reduced levels of Rav2 levels helps explain increased V-ATPase disassembly and addition of extra Rav2 improves assembly, we do not know whether the Rav2 reduction occurs as a result of some general vulnerability in aging cells or is deliberately programmed (Gladyshev et al., 2021). The significant increase in lifespan and prolonged V-ATPase assembly in *oxr1Δ* cells seems to argue against any growth advantage from increased V-ATPase disassembly in older cells, at least at the single cell level.

Many aging pathways are tightly linked to nutritional signaling, and reversible disassembly is often driven by nutritional signaling pathways. Given the rapid disassembly of the V-ATPase upon acute glucose deprivation (Kane, 1995), it was initially surprising that growth under CR conditions suppresses V-ATPase disassembly with age. However, we previously showed that the acute disassembly response required glucose concentrations well below 0.5% (Parra & Kane, 1998). Activation of the PKA pathway in the *ira2Δ* mutant (Supporting Information Figure 1b) promotes V-ATPase assembly in older cells, but also results in a decrease in RLS. In mammalian cells, V-ATPase assembly generally increases in response to nutrient deprivation or inhibition of mTOR function (Ratto et al., 2022). CR could mimic this effect in aging cells, preserving lysosomal acidification and function. Deletion of the mTOR effector *SCH9* extends RLS and CLS and promotes V-ATPase assembly (Wilms et al., 2017), similar to the effect of *oxr1Δ* mutation. Signals involved in reversible disassembly of the V-ATPase are incompletely understood and vary between physiological settings (Collins & Forgac, 2020). The RAVE complex responds to nutritional signals associated with reversible disassembly with release from the vacuolar membrane upon acute glucose deprivation and recruitment to the membrane upon glucose restoration, even in the absence of V1 subunit C and the V1 subcomplex (Jaskolka, Winkley, et al., 2021). The molecular mechanism of Oxr1-induced V-ATPase assembly is understood (Khan & Wilkens, 2024), but its response to cellular signaling is still being explored.

Reduced V-ATPase assembly could be a factor in the age-related decline in lysosomal acidification and function in higher eukaryotes. Reversible disassembly occurs in higher eukaryotes including mammalian cells. Rabconnectin-3 complexes of higher eukaryotes are functional homologues of the yeast RAVE complex (Jaskolka, Winkley, et al., 2021). Oxr1 belongs to a family of TLDc proteins and several of these proteins have been shown to bind to mammalian V-ATPases and induce disassembly (Oot & Wilkens, 2024). These data suggest that the core elements for controlled V-ATPase disassembly during aging are present in other cells. We observed V-ATPase disassembly in a yeast replicative aging model, which is most comparable to mammalian cell types that continue to divide, such as adult stem cells (He et al., 2018), and observed similar effects of *rav1Δ* and *oxr1Δ* mutations in chronological aging, which is more analogous to long-lived, non-dividing mammalian cells. Taken together, these data suggest that V-ATPase assembly state is linked to different types of aging and could easily play a role in aging in higher eukaryotes.

## METHODS

### Yeast strains and plasmids

All strains analyzed were in the BY4741 (*MATa his3*Δ*1 leu2*Δ*0 met15*Δ*0 ura3*Δ*0*) or BY4742 (*MATα his3*Δ*1 leu2*Δ*0 lys1Δ ura3*Δ*0*) background. BY4742 strains containing Vma5-GFP::*HIS3* and Vph1-GFP::*HIS3* were constructed as part of the genome-wide GFP-tagging project (Huh et al., 2003). The *rav1Δ::kanMX, oxr1Δ::kanMX, vma1Δ::kanMX*, and *ira2Δ::kanMX* mutant strains in the BY4741 background were purchased as part of a haploid deletion collection.

*VMA2* was C-terminally tagged with GFP by PCR amplification from pFA6a-GFP-KanMX6 and genomic integration (Longtine et al., 1998). The *oxr1Δ::kanMX* Vma5-GFP::*HIS3* strain was obtained by crossing BY4741 *oxr1Δ::kanMX* and BY4742 Vma5-GFP::*HIS3* and obtaining spores with the desired genotype by tetrad dissection. The *stv1Δ::natMX* mutation was introduced by crossing this strain to a *stv1Δ::natMX* haploid and sporulating. The pRS316 Oxr1- HA plasmid (Khan & Wilkens, 2024) was transformed into *oxr1Δ::kanMX* and transformants were selected on medium lacking uracil. Rav1-myc13::*kanMX*, Rav2-myc13::*kanMX*, and Rav2-FLAG::*kanMX* strains were constructed as described previously (Jaskolka, Tarsio, et al., 2021; Smardon et al., 2002). The tagged alleles were PCR amplified from the original strains and integrated into BY4741.

### Fluorescence microscopy

Strains expressing GFP-tagged Vma5, Vma2, or Vph1 were grown to log phase in synthetic complete (SC) medium containing either 2% or 0.5% glucose overnight, pelleted by centrifugation, then suspended in fresh medium containing 2% or 0.5% glucose and grown for an additional 2 hours. Cells were stained by addition of CW (final concentration 10 μg/ml) 5 min prior to imaging. Cells were visualized with a 100x oil (NA 1.4) objective on a Zeiss Imager.Z1 fluorescence microscope with a Hamamastu CCD camera and AxioVision software. Cells were viewed through DIC optics and fluorescence was imaged using a DAPI filter set (for CW) and a GFP filter set. Bud scars were counted from CW staining; in mixed age populations, cells with 5 or more bud scars were designated as old and those with less than 5 as young. To obtain a more precise count of bud scars for binning, cells were visualized on multiple focal planes. GFP fluorescence images were captured for cells in each binned range then processed in FIJI. To assess vacuolar localization, a line was drawn across a cell and through the vacuolar membrane. From this line scan, a plot illustrating fluorescence intensity along the line was generated, with peaks indicating areas of elevated fluorescence at the vacuolar membrane. The peak intensity (maximum fluorescence) was quantitated as a measure of vacuolar localization. Each biological replicate corresponds to a distinct culture of yeast cells. Maximum fluorescence was quantitated for at least 20 young and 20 old cells per biological replicate. In cells were binned by age, at least 20 cells per biological replicate were counted for each bin. Maximum fluorescence from line scans of young and old cells in each biological replicate were averaged and then divided by the average for young cells in that replicate to normalize. Normalized fluorescence for each biological replicate was plotted, along with average intensities across replicates +/- s.e.m. (standard error of the mean). Statistical significance was determined by t-test for the young and old cell comparisons and by ANOVA for binned samples. To show the range of values for the young cells across experiments, the values for young cells across biological replicates were averaged; individual replicates were divided by this average are shown as points on each graph.

### Age Enrichment

Age enrichment was performed as described by Jin et al. (Jin et al., 2021). Cells were collected from a 50 mL fresh overnight culture in YEP (yeast extract-peptone medium) supplemented with 2% or 0.5% glucose to an OD600 of 1.0 and washed twice with cold phosphate-buffered saline (PBS), pH 7.4. Cells were pelleted by centrifugation and washed three times in cold sterile PBS, then labeled with 1.6 mg/ml EZ-Link Sulfo-NHS-LC-Biotin (Pierce) at room temperature for 30 min with gentle agitation. After labeling, the cells were washed three times with cold PBS, pH 8.0, to remove free biotin, and resuspended in YEP medium supplemented with 2% or 0.5% glucose for growth overnight. After 16 hours, cells were pelleted by centrifugation and resuspended in 35 ml of cold PBS, pH 7.4, mixed with 250 μl of a magnetic streptavidin bead suspension (Pierce), and incubated for 60 min at 4°C. The mixture, in a 50 mL conical tube, was loaded onto a magnetic separation column (Permagen) at 4°C to separate biotinylated cells.

Unbound (young) cells were removed gently by pipetting. Magnetically separated cells were subsequently washed three times by resuspending in 35 mL PBS, pH 7.4 supplemented with 2% or 0.5% glucose, repeating magnetic separation, and discarding the supernatant. After washing, cells were resuspended in 200 mL of YEP or SC containing 2% or 0.5% glucose and allowed to grow for an additional 4 hours before obtaining the final “old” mother cells and “young” daughter cells. Cells were loaded on the magnetic separation column as described above and daughter cells obtained from the supernatant were pelleted by centrifugation to obtain a concentrated population of young cells. After the final wash, the magnetic beads were resuspended in 1 ml PBS, pH 7.4 and centrifuged to concentrate the old population. The two populations were then used immediately for pH measurements or stored at -80°C for further biochemical analysis.

### Whole Cell Lysates and Immunoblots

Cell pellets from age-enriched young and old cell populations were resuspended in cracking buffer (8 M urea, 5% SDS, 1 mM EDTA, 50 mM Tris-HCl, pH 6.8) at 95°C and lysed with glass beads. Cellular debris was pelleted by centrifugation at 16,000 × g for 2 min, and supernatants containing whole-cell extracts were used immediately or stored at −80 °C until use.

After determination of protein concentrations by Bradford assay, equal amounts of protein were separated by SDS-PAGE and transferred to a nitrocellulose membrane. Blots were blocked, then incubated overnight with primary antibodies at 4°C. Primary antibodies (all at a 1:500 dilution) included mouse monoclonal antibodies 7A2 against Vma5, 10D7 against Vph1, 8B1 against Vma1, and 13D11 against Vma2 (Kane, Kuehn, Howald-Stevenson, & Stevens, 1992). In addition, anti-myc monoclonal 9E10 (Santa Cruz Biotechnology), anti-HA monoclonal (BioLegend), anti-FLAG (Sigma) and anti-GAPDH (Proteintech) antibodies were used at 1:500, 1:500, 1:1000, and 1:10,000 dilution, respectively. HRP-conjugated anti-mouse secondary antibody (Bio-Rad) was added at a final dilution of 1:2000 and incubated for 60 min at room temperature. The blot was washed, incubated with Bio-Rad Clarity Western ECL substrate and imaged in an Azure Sapphire FL Biomolecular Imager. Images were quantified using FIJI.

Molecular mass markers were included on every blot. In blot images, the nearest molecular mass marker is indicated.

### Vacuolar pH Measurements

Vacuolar pH was measured using BCECF-AM (Invitrogen) as described previously (Diakov et al., 2013). Age-enriched populations of cells were loaded with BCECF-AM in YEP supplemented with 2% or 0.5% glucose. After washing with YEP media to remove dye, cells were resuspended in YEP, deprived of glucose, and incubated for 30 min. For fluorescence measurement, 20 μl of cell suspension was diluted into 3 ml 1 mM 2-(N-morpholino)ethanesulfonic acid (MES) pH 5.0 buffer, and fluorescence intensity at excitation wavelengths 450 and 495 nm and emission wavelength 535 nm and was monitored continuously in a Horiba Jovin Yvon Spectrafluor Max fluorometer at 30°C. The fluorescence ratio for each sample was calibrated in every experiment by clamping pH in the range of 5.0 to 7.0 and constructing a calibration curve.

### Replicative Life Span

RLS was determined on YEP, 2% glucose plates buffered to pH 5. Daughter cells were sequentially removed by micromanipulation (Steffen, Kennedy, & Kaeberlein, 2009). Survival curves are pooled data from experiment-matched controls. The number of divisions for 26-30 mother cells were scored for each curve. p-values for replicative life span survival curve comparisons were calculated with a Cox regression model. Kaplan-Meier survival curves were plotted with GraphPad Prism.

### Chronological lifespan analysis

CLS was assessed as described (Wilms et al., 2017), with the following modifications. Cells were pre-cultured to the stationary phase in SC containing 2% glucose. Stationary-phase cells were then diluted to an OD600 of 0.1 in fresh medium and grown for 48 hours; the end of this time is designated as time zero. Cells were aged in SC containing 2% glucose adjusted to pH 5.5 with 500 mM MES. Cell viability and clonogenicity (colony forming units; CFUs) were analyzed on days 7, 14, 21, 28, and 35. Cell death was assessed by propidium iodide staining (Sigma) and visualized by fluorescence microscopy. For CFU analysis, cell density was measured using a hemocytometer, and 250 cells were plated onto YPD plates. After incubation, CFUs were counted using FIJI, and the results expressed as the percentage of total cells plated. Statistical significance was assessed by ANOVA with multiple sequence comparisons (GraphPad Prism).

### RT-PCR of RAV2 from young and old cells

RNA was extracted from young and old cells, obtained by age enrichment as described above, using the NEB Monarch Total RNA Miniprep Kit. RT-PCR was conducted using the NEB Luna Universal One-Step RT-qPCR Kit and performed on a Bio-Rad CFX384 Touch System. Data analysis was conducted using CFX Maestro Software.

## Acknowledgements

This work was supported by NIH R35 GM145256 to P.M.K. The authors thank M. Murad Khan and Dr. Stephan Wilkens for sharing the *oxr1Δ* strain and Oxr1-HA plasmid, and Dr. Xin Jie Chen for helpful discussions and a critical reading of the manuscript. Illustrations in Figures 2a and 5a were prepared using Biorender.com.

## Author contributions

F.H. performed experiments, analyzed data, prepared figures, and wrote the first draft of the manuscript; P.M.K. obtained funding for the project, analyzed data, prepared figures, and contributed to writing of the manuscript.

## Data availability

The data that support the findings of this study are available at Upstate.figshare.com. https://upstate.figshare.com. DOI: 10.58120/upstate.27902313

