## Supporting Information Figure 1 for "V-ATPase Disassembly at the Yeast Lysosome-Like Vacuole Is a Phenotypic Driver of Lysosome Dysfunction in Replicative Aging"

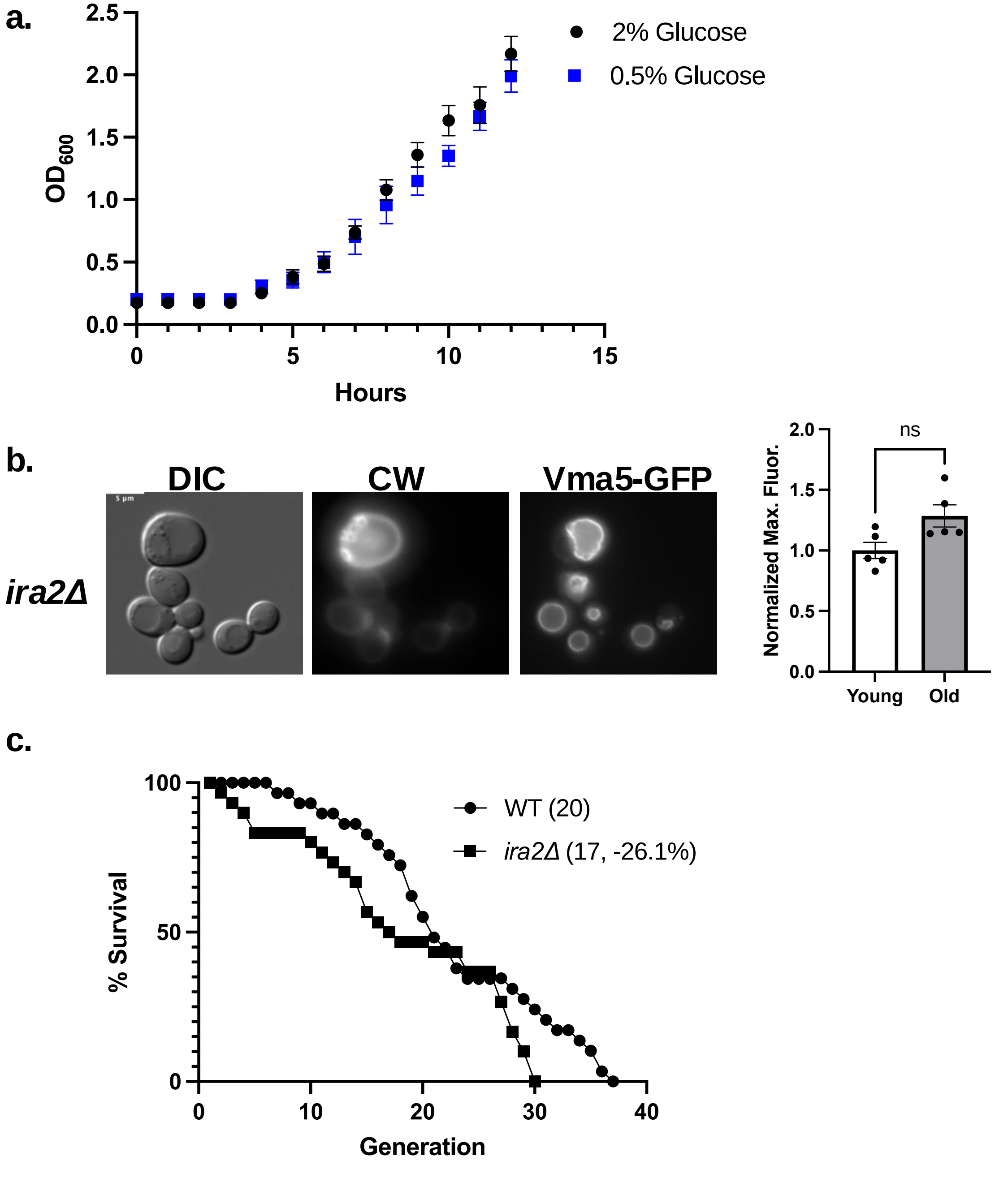


**Supporting information, Figure 1: a.** Cells show similar growth rates over 12 hours in 2% and 0.5% glucose. BY4742 cells at log phase were diluted into YEP containing either 2% or 0.5% glucose as indicated. $OD_{600}$ was measured every hour for 12 hours. b. BY4742 *ira2*Δ cells expressing Vma5-GFP grow in SC with 2% glucose. CW was used to visualize bud scars. Normalized maximum fluorescence was obtained through line scan quantitation using FIJI as in **Figure 1**. Means +/- s.e.m. of five biological replicates are shown; each replicate is represented by a dot. Significance was calculated by unpaired Student's *t* test and results were not significant (n.s.). c. Kaplan-Meier curves comparing the RLS of *ira2*Δ (squares), and wild-type cells (circles). 29 wild-type cells were scored and 30 *ira2*Δ. Deletion of *IRA2* significantly shortens RLS ($p < 0.01$).
