## Supporting Information Figure 2 for "V-ATPase Disassembly at the Yeast Lysosome-Like Vacuole Is a Phenotypic Driver of Lysosome Dysfunction in Replicative Aging"

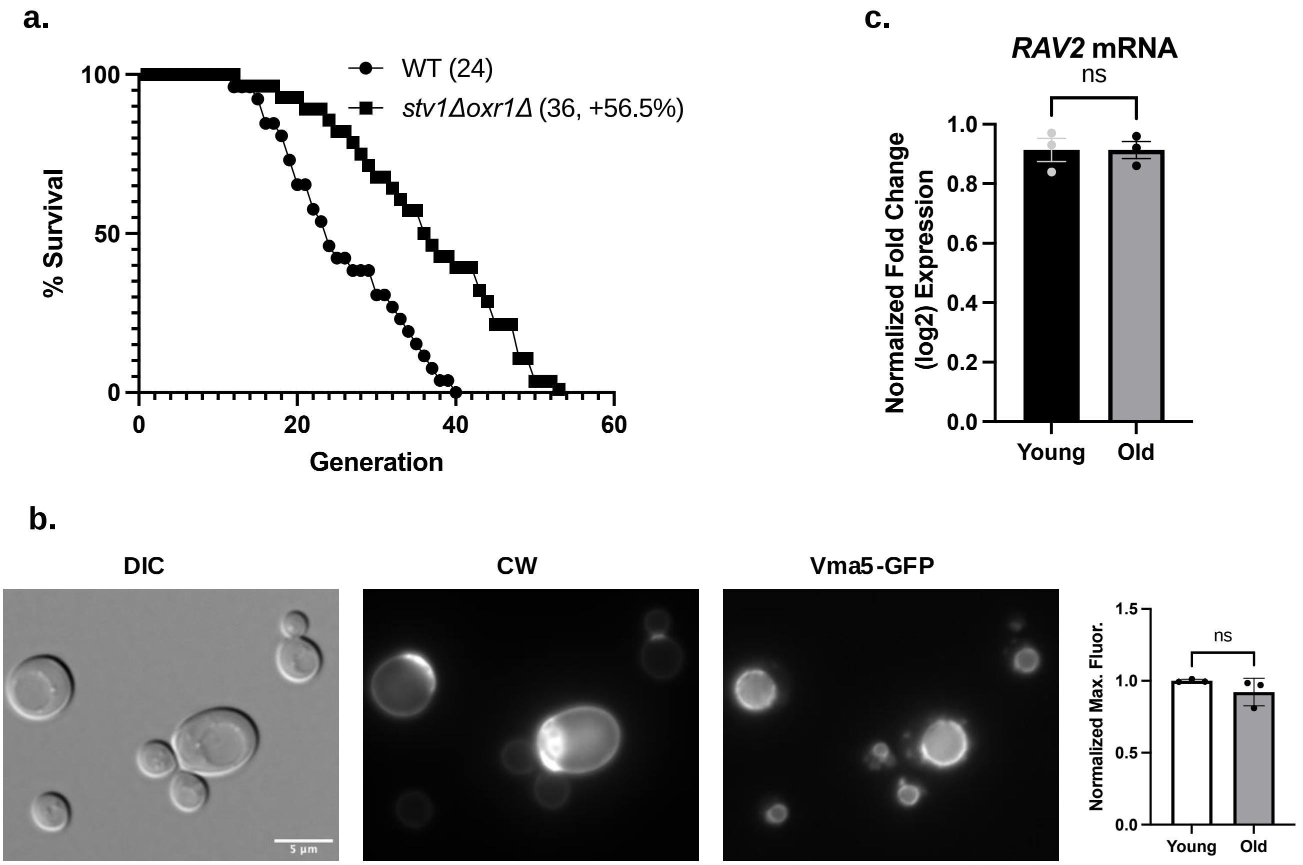


**Supporting information Figure 2:** a. Kaplan-Meier curves comparing the RLS of *stv1*Δ *oxr1*Δ (squares), and wild-type cells (circles). 26 wild-type cells were scored and 28 *stv1*Δ *oxr1*Δ. $p < 0.001$. b. BY4742 *stv1*Δ *oxr1*Δ cells expressing Vma5-GFP grow in SC with 2% glucose. CW was used to visualize bud scars. Normalized maximum fluorescence was obtained through line scan quantitation of Vma5-GFP using FIJI as in **Figure 1**. Means +/- s.e.m. of three biological replicates are shown; each replicate is represented by a dot. Significance was calculated by unpaired Student's *t* test and the difference was not significant (n.s.). c. Quantitative RT-PCR comparing expression of *RAV2* mRNA between young and old cells; the statistical difference was not significant (n.s.).
